# The Effect of Visual Impairment on the Controllability of a Brain-Computer Interface Using Visual Spatial Attention

**DOI:** 10.64898/2026.09.01.748503

**Authors:** Christoph Reichert, Mircea Ariel Schoenfeld, Stefan Dürschmid

## Abstract

The primary goal of a brain-computer interface (BCI) is to enable interactions with the environment by generating control signals directly from brain activity, independent of physical movement including gaze shifts. Attentional selection of peripheral objects has been used to control a BCI but requires sufficient visual acuity to reliably recognize the target stimulus. Here, we systematically investigated the effect of different stages of visual impairment on event-related potentials (ERPs) and BCI performance while healthy participants directed covert attention to peripheral visual stimuli. We hypothesized that the effect of visual impairments such as blurred or monocular vision might alter attention-related visual ERPs, but has only minor or no impact at all on BCI performance when color is the target feature. Early attention-sensitive visual components were altered in monocular vision compared to binocular vision. Simulated vision degradation resulted in reduction of ERPs in the P300 range. The differences in ERPs across vision conditions did not affect the decoding accuracy of the BCI. The study not only revealed differences of visual ERPs in different viewing conditions but primarily demonstrated that a BCI can be reliably controlled by focusing attention on a visual stimulus whose distinguishing features such as color, can be recognized even with visual impairments. This provides a promising approach to enable paralyzed persons to communicate using a non-invasive, gaze-independent BCI.

## 1. Introduction

Visual processing in the human brain is commonly investigated in healthy participants with normal or corrected-to normal vision. However, BCI approaches that rely on precise visual perception may have limited practical utility because many potential BCI users often suffer from visual impairments due to age, insufficiently corrected ametropia, or paralysis of extraocular muscles. The inability of shifting attention to a target stimulus diminishes the amplitude of brain potentials related to spatial attention. As a consequence, BCI applications that proved to be accurate in healthy participants might be difficult to control by the potential end-users.

A prominent example of an BCI that relies on intact vision is the visual P300 speller. This BCI identifies intended letters from event-related P300 responses, evoked when an attended target letter in a matrix of letters is flashed (Fazel-Rezai et al., 2012). McCane et al. examined the effects of amyotrophic lateral sclerosis (ALS) on BCI performance and found that while the decoding accuracy remained comparable to that of healthy controls, the associated event-related potential patterns differed significantly (McCane et al., 2015). Importantly, in this study the patients had no major visual impairments. However, in an earlier study McCane et al. also reported that a group of patients suffering from vision impairment could not gain BCI control (McCane et al., 2014). To overcome limitations associated with impaired vision, auditory and tactile stimuli have been used to evoke P300 potentials for BCI control in paralysed persons. However, these studies showed mixed results. Kübler et al. showed that auditory P300 performance is less reliable than visual P300 speller control (Kübler et al., 2009). Accordingly, a case study of a patient with locked-in syndrome revealed that tactile stimulation outperformed visual stimulation, while auditory stimulation produced lowest performance (Kaufmann et al., 2013). In contrast, in a cohort of eleven healthy participants, visual stimuli resulted in significantly higher accuracies than tactile and auditory stimuli (Halder et al., 2018). Together, these studies suggest that visual impairment is a critical factor in BCIs for patients.

Another class of BCIs is based on sensorimotor rhythms (SMRs), which are voluntarily modulated through motor imagery to enable device control. In motor imagery users train to modulate their SMR by imagining to move different limbs without any external stimuli to induce control signals. However, compared to P300 spellers, SMR requires a considerably higher amount of training sessions to achieve a necessary level to control a BCI and permits lower information transfer rates in ALS patients (Nijboer et al., 2010). While decoding of motor activity is a promising tool for BCI control, especially with implantable BCIs (Vansteensel et al., 2023), long-term stability of the neural signal is a limiting factor. Progressive reductions in neural signal amplitudes associated with disease progression can impair BCI performance, thereby limiting effective communication (Vansteensel et al., 2024, 2016).

Independence of eye movement is a prerequisite for patients with impaired oculomotor function. While motor imagery is intrinsically suited for gaze-independent BCI control, the matrix speller and most BCIs using visual stimuli to evoke visual potentials are not. Using rapid serial visual presentation of stimuli at the center of the visual field is one of the approaches to overcome this problem (Acqualagna and Blankertz, 2013; Lees et al., 2018), but the need to recognize the rapidly presented target items could pose a challenge for people with visual or cognitive impairments. Finally, visual spatial attention is a promising cognitive task that has been shown to enable reliable control of a binary communication system by decoding the shift of attention to the left or right visual field (Reichert et al., 2022, 2020b). Since there are only two alternatives to choose from, which can be selected by paying attention to one of two colors, this paradigm appears appropriate for people with visual impairments. Here we investigate whether the brain potentials elicited by attention to a target in the left and right visual field are preserved when vision is either blurred or one eye is blinded. We are particularly interested in whether and how the BCI performance might be affected in these conditions. Two experiments, each investigating three different conditions, were conducted to address this question.

## 2. Methods

### 2.1. Subjects and Task

Eighteen healthy participants (9 female), mean age 24.7 year (2.9) participated in two experiments which were conducted on two separate days. Participants gave written informed consent and received financial compensation for participation. For each participant, we tested their visual acuity using the Snellen chart and determined which eye was the dominant eye.

In two experiments we investigated the impact of vision impairment on the ability to control a brain-computer interface with feature-based attention to peripherally presented visual stimuli. Task and stimuli were the same in both experiments. To mimic potential visual impairments, we manipulated the normal vision of healthy participants in two ways. In experiment 1, we covered one eye, which could prevent diplopia resulting from impaired or paralyzed extraocular muscles, disrupting binocular alignment. In experiment 2, participants wore plano lens spectacles equipped with Bangerter occlusion foil, which are usually used in amblyopia therapy. The order of the two experiments was counterbalanced across participants.

We employed a BCI protocol based on visual spatial attention (VSA) that enables reliable control in two- and four-choice applications (Reichert et al., 2022, 2020a). We used a custom-made stimulation device with two quadratic LED panels spanning 2.86° visual angle (va) located 8.93° to the left and right relative to the fixation cross. The fixation cross, spanning 2.45°, was presented 2.86° above the panels to induce robust ERP components reflecting visual spatial attention (Luck et al., 1997). Behind the opal panes, we placed two letters (height 1.80° va) made of color filter foil: a red ‘Y’ and a green ‘N’. Consequently, when the panel was illuminated with green light, the ‘Y’ appeared darker and was more salient, and when illuminated with red light, the ‘N’ was more salient, since it appeared darker (see Figure 1). While the shape of the letter served as additional target feature for the visual search task, recognizing orthographic stimuli also requires additional cognitive resources (Winsler and Luck, 2026). For experimental control as well as online and offline data analysis we used MATLAB R2021b. LEDs were controlled using an Arduino Leonardo microcontroller (Arduino, Somerville, MA).

**Figure 1:**
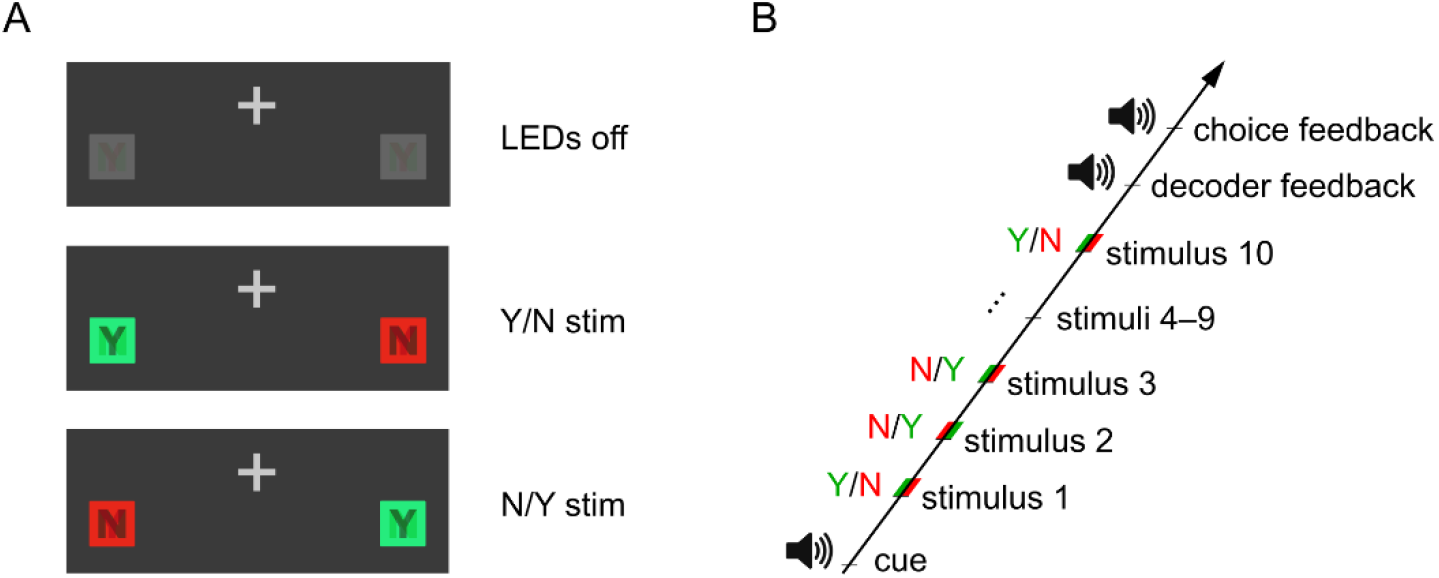
Stimulus and trial structure. **(A)** The stimulus board could have three states: Off, left LED emitted green light and right LED red light and vice versa. Depending on light color, the letter Y or N was more salient. **(B)** Cue, stimuli and feedback. After the cue was auditorily presented, ten visual stimuli were presented in random order. The prediction of the BCI and the correctness of the alleged choice was auditorily presented.

Participants were seated in an EEG cabin with the light dimmed. EEG was recorded at 500Hz from 29 Ag/AgCl electrodes, where placement followed the international 10-20 system, with additional electrodes placed at selected intermediate 10–10 locations, using a BrainAmp DC amplifier (Brain Products GmbH, Gilching, Germany) and referenced against the right mastoid. Simultaneously, the horizontal and vertical electrooculogram (EOG) was recorded. For online analysis, 14 parieto-occipital channels primarily involved in VSA processes were used. For offline analysis, 27 electrodes excluding T7 and T8, which showed muscle artefacts in five participants, were used.

Participants were asked to fixate the fixation cross while they were presented with a synthetic voice reading a random number between zero and ten. For each presented number (1-10) they answered the question “Was the number even?” by a covert shift of attention to the green (‘Yes’) or red stimulus (‘No’). If the number was zero, participants were asked to ignore both red and green stimuli, which enabled a contrast between attention and passive viewing. Following the presentation of the number plus a preparation interval (2.5 sec), pairs of red and green stimuli were repeatedly presented for approximately 10 s. Both stimuli appeared simultaneously for 250 ms, with the red on the left and the green stimulus on the right or vice versa, in randomized order. Consecutive presentations were separated by a stimulus onset asynchrony (SOA) of 700 ms plus a random jitter of 0-100 ms. Two seconds after the last stimulus presentation, the EEG signals of the entire presentation sequence were retrieved and processed. We decoded which stimulus participants attended to and, thus, which response they intended to communicate (see *Analysis of EEG data*). First, the decoded answer was presented by a synthetic voice saying ‘Yes’ or ‘No’. Second, auditory feedback indicated whether decoded and actual correct answer matched. Correct responses were indicated by two consecutive tones increasing in frequency (synthesized piano keys A4 and E5), whereas mismatches were signaled by two consecutive tones decreasing in frequency (synthesized piano keys A4 and D4). No feedback was provided in the passive viewing condition.

In both experiments we investigated three different viewing conditions, separated in blocks counterbalanced across subjects. Each block consisted of one training run where only the cues but no feedback was presented and three feedback runs. Each run consisted of 12 trials, with cues 1-10 presented once and cue 0 two times. The decoder model was first trained using data of the first run per block and retrained after each trial in the following runs. In experiment 1 viewing conditions were binocular vision (no eye covered), monocular right-eye vision (left eye covered) and monocular left-eye vision (right eye covered). In experiment 2 viewing conditions were normal vision (no spectacles), moderately degraded vision (Bangerter foil value 0.3), and highly degraded vision (Bangerter foil value 0.1), which simulate moderate and severe vision impairment (World Health Organization, 2019), respectively.

### 2.2. Analysis of EEG data

For both online and offline decoding of spatial attention we used the ERPCCA toolbox, which is designed to predict BCI commands from a series of event-related potentials (ERPs) (Reichert et al., 2024). The toolbox trains a model which estimates canonical components from brain signals and model signals by maximizing its correlation. The model signals provide the information needed to discriminate target from nontarget events. To decode the target from unseen data, the canonical coefficients obtained from the training data are applied to the test data and the resulting canonical components are used for classification.

We pre-processed the data for offline analysis as follows. First, we removed all inter-trial data and filtered the EEG and EOG signals between 0.25 Hz and 30 Hz using a 6th order Butterworth band-pass filter. We performed independent component analysis (ICA) using the FastICA toolbox (Hyvärinen and Oja, 2000) to remove activity related to eye movements. Specifically, we calculated the L2 norm of correlation coefficients between a given independent component (IC) and both the horizontal and vertical EOG. ICs exceeding a threshold of 0.5 were removed from the raw data using the mixing and demixing matrices obtained by ICA on the filtered data. Afterwards, we re-referenced the EEG data to the mean of left and right mastoid. Line noise was removed by a 50 Hz notch filter. Artefact detection was performed by determining the envelope of the band-pass filtered EEG signals (1 Hz–30 Hz) by Hilbert transform and labelling an epoch as artefactual if any value exceeded the threshold 50 µV. For decoding, the brain signals were filtered between 1 Hz and 12.5 Hz and resampled to 50 Hz to reduce the matrix sizes for the CCA. For statistical analysis, the resulting time series were filtered between 0.1 Hz and 25 Hz and resampled to 125 Hz. We then defined stimulus epochs between 0 and 600 ms relative to stimulus onset for decoding. For statistical analysis stimulus epochs were defined from 0 to 750ms, which were corrected for baseline activity (–200 to 0 ms). For online decoding, we performed the same pre-processing steps but did not perform ICA and artefact detection.

In order to decode the target color to which attention was shifted, we selected the parameters for the CCA-based model estimation as follows, motivated by results of recent studies of our group (Reichert et al., 2024). To isolate attentional allocation reflected in difference waves, we used impulse functions as model signals and enabled the contrast parameter. The number of canonical components used for classification was limited to 3 and class sizes were set to be balanced in training sets. As feature space we selected the correlation values which internally were used to classify the target by determining the maximum correlation in a stimulus sequence, which was comprised of the ten epochs in a trial.

In order to decode *attention* and *passive viewing*, we applied the same pre-processing steps and model parameters as in decoding the target color but provided different condition labels accordingly. Specifically, all epochs comprising an *attention* trial regardless of the target color were labelled as the first condition and all epochs comprising a *passive viewing* trial were labelled as the second condition. To predict whether the stimuli in a trial were ignored or not all epochs comprising a trial were applied to the decoder model.

### 2.3. Analysis of EOG data

First, we investigated to what extent EOG could be used to decode attentional shifts. We trained a decoder model with the EOG data and determined decoding accuracies as outlined above. Second, we calculated the deflection of the horizontal EOG as an estimate of involuntary eye movements in different viewing conditions. Specifically, we calculated the average of the five highest absolute values, each of which covers the sampling interval of 20 ms.

### 2.4. Validation and statistical analyses

All offline decoding results we report are based on leave-one-out cross-validation (CV), i.e., each trial in a subset of interest is decoded once by using all other trials of the subset as training trials. To determine statistical significance, we estimated the confidence level around the empirical chance level by permutation tests. Specifically, we determined a distribution of decoding accuracies obtained by permuting the labels of the target sides and performing a CV with this brain data set uncorrelated to the task. This procedure was repeated 1000 times and the 97.5th percentile of the performance measures served as upper chance level, i.e. the upper bound of the 95 % confidence interval.

We performed nonparametric statistical tests to determine significant differences. Specifically, we performed Wilcoxon signed-rank tests to compare two conditions and report the p-value as p_sr_. To derive statistical inferences of the ERP data between viewing conditions, we performed cluster-based permutation tests (Maris and Oostenveld, 2007). The advantage of this approach is that in a data set with spatially and temporally correlated samples the challenge of multiple comparisons is effectively controlled. Specifically, we used Wilcoxon signed-rank tests to derive the actual test statistic as a matrix of z-values. We determined clusters of significant z-values and summed z-values within a cluster as z_clust_, denoting the cluster mass. Then we randomly drew samples from the two test sets in new test sets of the same size and repeated the signed-rank test 10,000 times. Clusters with the largest cluster mass were pooled from each repetition to create a null distribution of z_clust_ values. Finally, the actual cluster mass is compared to the null distribution to obtain a p-value that indicates the significance of the actual cluster.

## 3. Results

We estimated the participant’s initial visual acuity by presenting a Snellen chart at the same distance as the stimuli were presented (70 cm). Visual acuity was ≥1.0 in 15 participants, ≥0.8 in two participants, and 0.67 in only one participant. In 13 participants the right eye was the dominant eye. We presented auditory feedback derived from the cue and the colour that was decoded as target colour starting after one calibration run per condition. The low number of training data resulted in significantly lower online decoding results as compared to a cross-validation (CV) approach. In experiment 1, the average online decoding accuracy of detecting the target, to which attention was shifted, was µ=82.6% (σ=11.9%) in the binocular vision (BV) condition, µ=78.2% (σ=12.4%) in the monocular right-eye vision (MREV) condition, and µ=77.8% (σ=13.0%) in the monocular left-eye vision (MLEV) condition. In experiment 2 it was µ=79.6% (σ=14.4%) in the normal vision (NV, no foil) condition, µ=78.7% (σ=12.3%) in the moderately degraded vision condition (MDV, foil value .3), and µ=73.9% (σ=13.7%) in the highly degraded vision (HDV, foil value .1) condition. Statistical analyses were performed on decoding accuracies obtained by leave-one-out CV and will be reported below. CV benefits from a greater number of training samples per condition providing a more reliable estimate of the performance of a sufficiently trained decoding model.

### 3.1. Experiment 1: Monocular vision and binocular vision

#### 3.1.1. Effects of vision impairment

We used cluster-based permutation tests and compared the EEG signals following stimulus onset to determine differences between binocular and monocular vision. A cluster of 16 parieto-occipital electrode sites (z_clust_=-369, p_clust_=0.0320) showed a stronger deflection to BV vs. MREV between 132ms and 164ms following stimulus onset (peak difference at 148ms at electrode PO4; p_sr_ =.0002, see Figure 2A). In this interval, BV led to an earlier rise of the N1 component with higher amplitude compared to MREV. Similarly, a second cluster indicates a stronger rise in component P2, which results in a higher amplitude during BV. Here, the maximum difference was at 205ms at Pz (p_sr_ =.0009), where it ranged from 181ms to 222ms and involved 16 electrodes as well (z_clust_=321, p_clust_=0.0460). The same pattern of clusters was found when we compared BV with MLEV but with the first cluster located towards the left hemisphere (Figure 2B). Specifically, the greatest difference was at 140ms at electrode P3 (p_sr_ =.0005). Here, the temporal range of this cluster was from 123ms to 156ms involving 15 electrode sites (z_clust_=-299, p_clust_=0.0589). The difference wave of the second cluster peaked at 197ms at Pz as well (p_sr_ =.0025), where the time range was identical to that of MREV and 19 electrode sites were involved (z_clust_=347, p_clust_=0.0450). No significant cluster for a difference between MREV and MLEV has been found. Including all trials, attention and passive viewing trials, the cluster-based permutation test revealed early clusters for BV vs. MREV (z_clust_=140, p_clust_=0.1988) and BV vs. MLEV (z_clust_=183, p_clust_=0.1648), which did not meet the significance criterion of p_clust_ <0.05, but the p-value at the difference peak (MREV: 99ms at PO4, p_sr_=0.0002; MLEV: 107ms at PO4, p_sr_=0.0006) clearly indicated a significant difference. Similar to the other effects, showing the highest difference at the rising flank, the P1 component rises earlier and stronger in bilateral compared to monocular vision.

**Figure 2:**
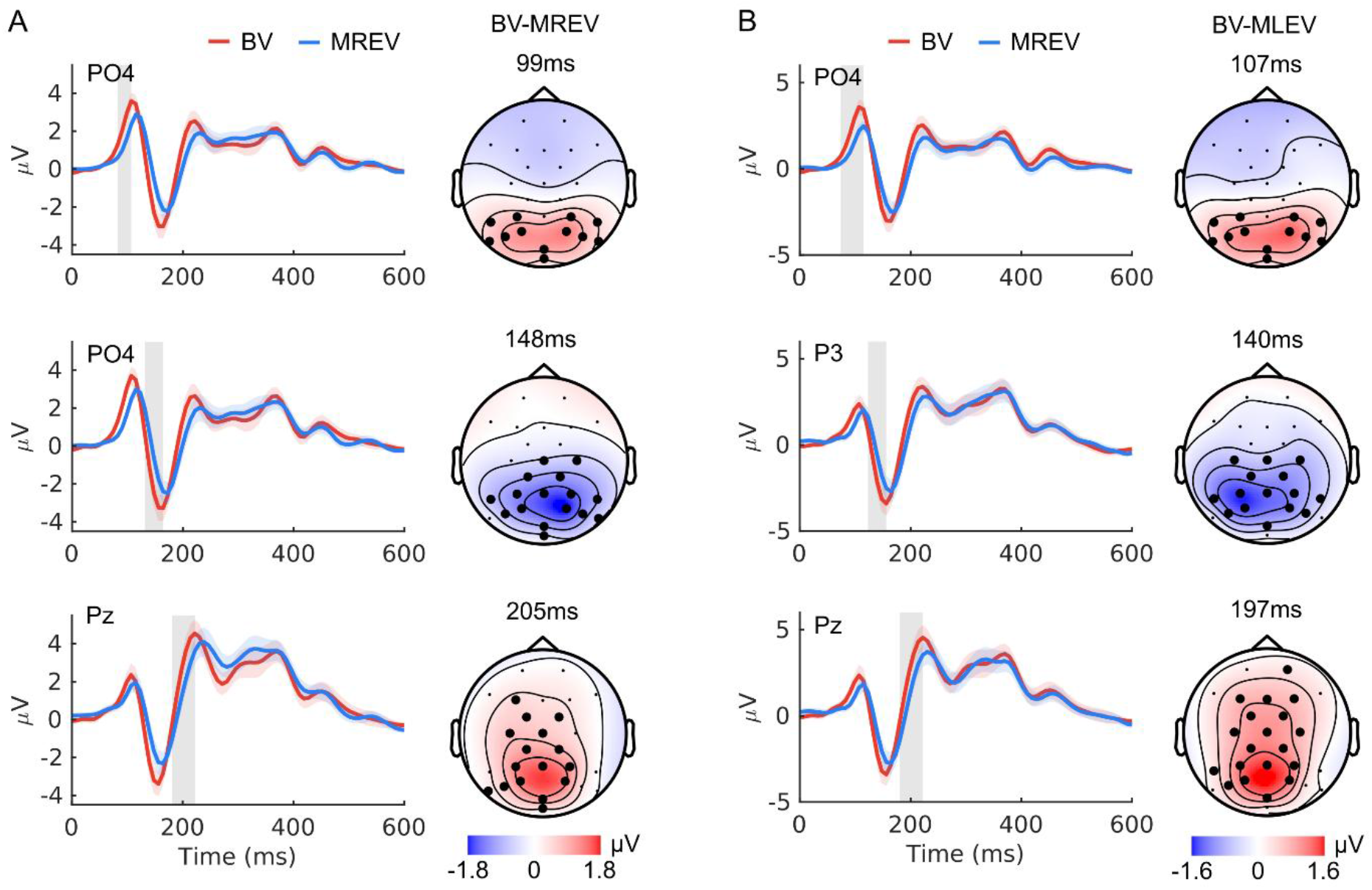
Monocular vision effects. Diagrams show the average signals per condition at the electrode with greatest difference in a cluster. Grey areas indicate the range of cluster time points at that electrode. Enlarged electrode markers indicate cluster channels at the time point of greatest difference. **(A)** Binocular vision (BV) compared to monocular right-eye vision (MREV) revealed clusters at the increasing flanks of P1, N2 and P2. **(B)** BV compared to monocular left-eye vision (MLEV) induced similar clusters as in MREV but with an apparent laterality shift in the N1 cluster.

#### 3.1.2. Decoding accuracy

We compared decoding accuracies of laterally presented targets using CV within and across conditions. In the BV runs the average decoding accuracy was µ=93.7% (σ=6.4%), while with MREV the CV revealed an accuracy of µ=92.1% (σ=8.0%) and with MLEV accuracy was µ=93.1% (σ=6.4%), showing no significant difference between conditions (see Figure 3). The absence of significant differences in decoding accuracies suggests that brain processes related to the attention task are similar across conditions, which is also supported by the high decoding accuracy achieved using all trials of the experiment (µ=94.4%, σ=4.7%), demonstrating generalization across conditions. Generalization is also indicated by the results shown in Table 1, where decoding accuracies were obtained using training data of one condition and predicting the target using only the trials of another condition. Only when the decoding model was trained on trials from the MREV condition and tested on trials from the MLEV condition, the accuracy decreased significantly compared to the within-condition CV of the MLEV condition (p_sr_=0.019).

**Figure 3:**
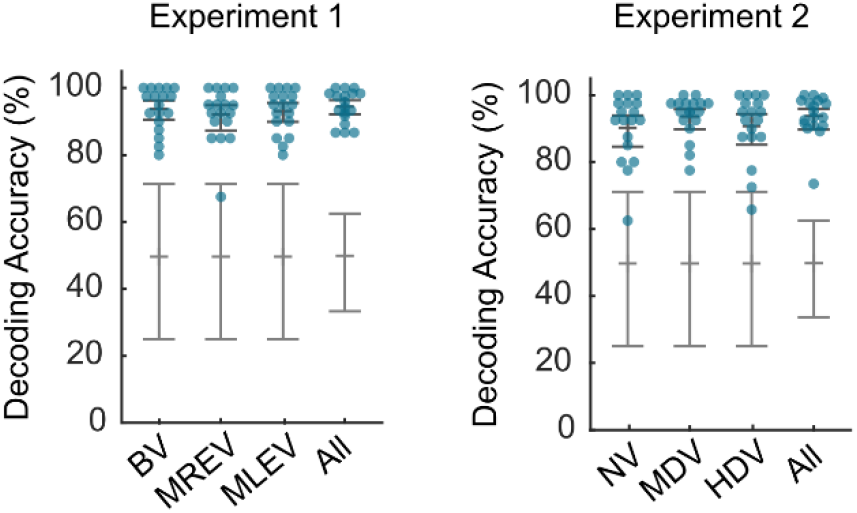
Decoding results. Decoding accuracies obtained by leave-one-out CV in both experiments. Blue circles indicate single subject performance and error bars indicate the 95% confidence interval of the mean obtained by bootstrapping. Grey error bars indicate the chance distribution obtained by permutation testing (95% percentile).

**Table 1:** Generalization across conditions in experiment 1. Average decoding accuracy is given in % correct and standard deviation in parentheses. Asterisks indicate significant difference from the corresponding condition’s within-condition CV accuracy with p<0.05.

|  | test trials |  |  |  |
| --- | --- | --- | --- | --- |
|  | vision | BV | MREV | MLEV |
| training trials | BV | - | 89.0 (8.7) | 90.3 (9.1) |
|  | MREV | 92.1 (6.3) | - | 88.9 (8.7)* |
|  | MLEV | 90.7 (8.4) | 89.9 (6.6) | - |

### 3.2. Experiment 2: Visual acuity

#### 3.2.1. Effects of vision impairment

The visual acuity conditions revealed no significant cluster when we compared NV and MDV. However, a cluster of 19 electrode sites (z_clust_=586, p_clust_=0.0070) showed a more positive wave in NV compared to HDV between 321ms and 370ms following stimulus onset (peak difference at 345ms at PO4, p_sr_ =.0043, see Figure 4A) and exhibited a bilateral parietal pattern. Comparing MDV with HDV revealed a cluster of 16 electrode sites (z_clust_=388, p_clust_=0.0140) with a more central spatial distribution, showing a stronger deflection in MDV vs. HDV between 321ms to 370ms following stimulus onset (peak difference at 337ms at electrode Cz; p_sr_ =.0043; see Figure 4B). In contrast to Experiment 1, including all trials, attention and passive viewing trials, neither a cluster nor a significant difference in single signed-rank tests were found in the P1 range.

**Figure 4:**
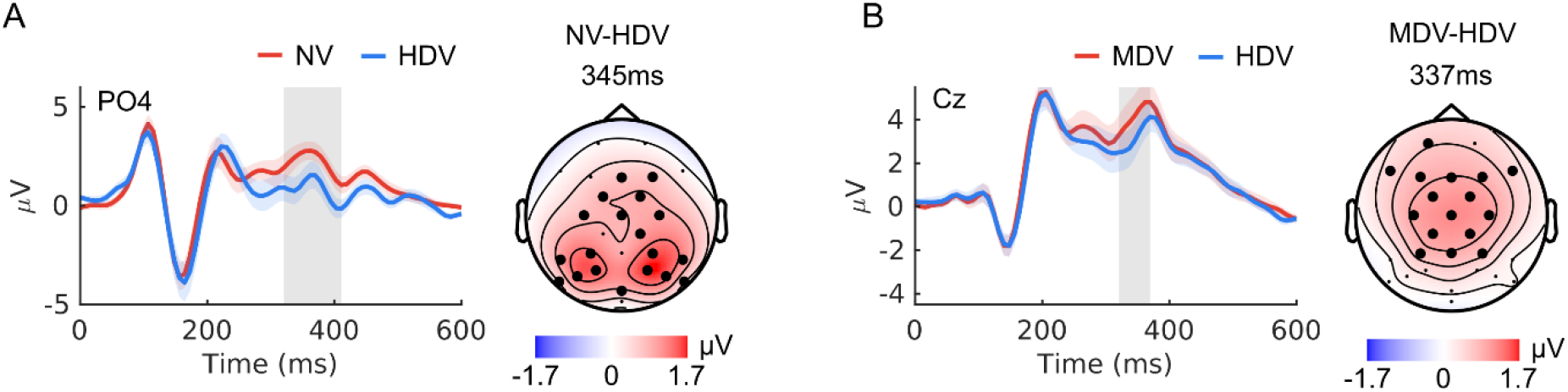
Blurred vision effects. Diagrams show the average signals per condition at the electrode with greatest difference in a cluster. Grey areas indicate the range of cluster time points at that electrode. Enlarged electrode markers indicate cluster channels at the time point of greatest difference. **(A)** Normal vision (NV) compared to high degraded vision (HDV) induced a greater positive response in a cluster around 350ms with bilateral parieto-occipital distribution. **(B)** Moderately degraded vision (MDV) compared to HDV induced a cluster in the same time range but with a central distribution.

#### 3.2.2. Decoding accuracy

In Experiment 2, the NV condition resulted in an average decoding accuracy of µ=90.3% (σ=10.0%). MDV resulted in not significantly different decoding accuracies of µ=93.6% (σ=6.2%) and HDV in µ=90.7% (σ=9.9%), showing no significant difference to the other conditions (see Figure 3 for details). Using all trials of the experiment resulted in µ=93.8% (σ=6.2%), indicating generalizability across conditions, similarly as observed in experiment 1. When we trained the decoding model with trials of one condition and the test set consisted of trials from the other conditions, generalizability was reduced compared to experiment 1 but still resulting in highly reliable accuracies (see Table 2). Significantly reduced accuracies compared to within-condition CVs were obtained when training on HDV trials and testing on MDV trials (p_sr_=0.005) and when training on NV trials and testing on MDV trials (p_sr_<0.001).

**Table 2:** Generalization across conditions in experiment 2. Average decoding accuracy is given in % correct and standard deviation in parentheses. Asterisks indicate significant difference from the corresponding condition’s within-condition CV accuracy with p<0.05.

|  | test trials |  |  |  |
| --- | --- | --- | --- | --- |
|  | vision | NV | MDV | HDV |
| training trials | NV | - | 88.7 (7.9)* | 87.2 (7.6) |
|  | MDV | 87.9 (10.2) | - | 88.8 (7.5) |
|  | HDV | 87.1 (10.0) | 89.5 (8.5)* | - |

### 3.3. Attention and passive viewing

#### 3.3.1. Effects of attention

In both experiments we used a cue (the number zero) that instructed the participants to ignore the stimulus without shifting attention to any of the visual stimuli. A clear difference in the average of EEG signals was evident between ignored trials and trials in which a target was selected. In experiment 1, a cluster-based permutation test revealed a significant cluster with maximum difference at 378ms after stimulus onset at electrode Cz (p_sr_ =.0002), where the *attention* condition resulted in a positive deflection compared to *passive viewing*. At that time point it extended across all electrodes except PO9, Iz and PO10. At Cz it ranged from 247ms to 485ms (z_clust_=2397, p_clust_=0.001) (Figure 5). A similar cluster was found in experiment 2, where the difference peak at Cz was at 370ms (p_sr_ =.0002) and the temporal extension of the cluster ranged from 263ms to 485ms (z_clust_=1929, p_clust_=0.001).

**Figure 5:**
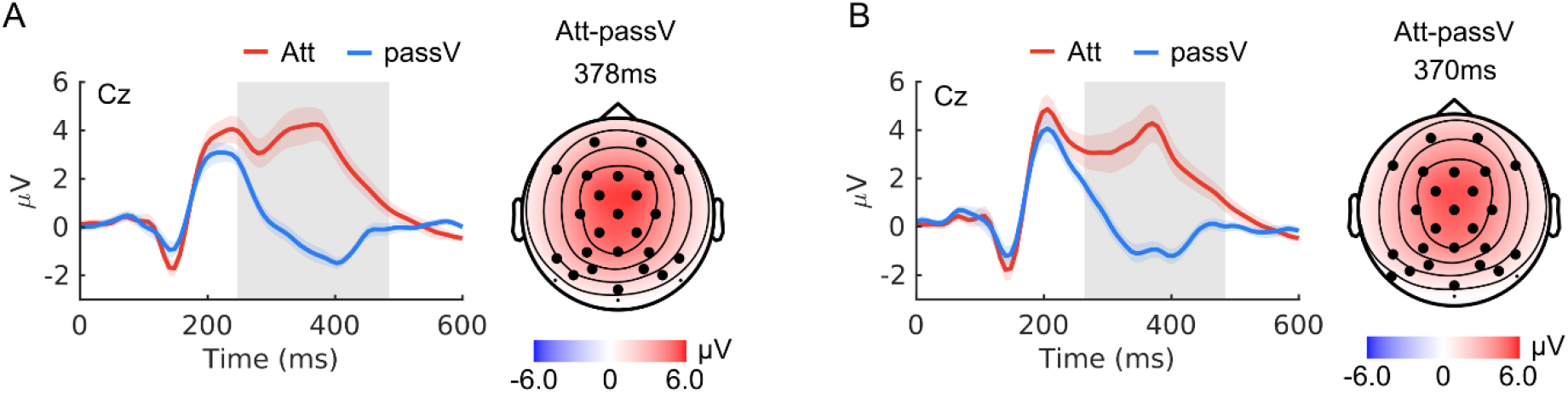
Attention effects. Diagrams show the average signals for the attention (Att) and passive viewing (passV) condition at the electrode with greatest difference in a cluster. Grey areas indicate the range of significant cluster time points at that electrode. Enlarged electrode markers indicate cluster channels at the time point of greatest difference. **(A)** P300 signal evoked by attention to a target in experiment 1 and **(B)** experiment 2.

#### 3.3.2. Decoding accuracy

We did not use the passive viewing data for online decoding during the experiment. In offline analyses these trials were used to distinguish between *attention* and *passive viewing* which ultimately indicates the participant’s intention for selection. Since we have only two of those trials per run compared to ten attention trials per run, we only report decoding accuracies using all trials to rely on sufficient training data. Additionally, we report sensitivity and specificity of attention detection to take a potential bias caused by the class imbalance into account. In experiment 1, we achieved a decoding accuracy of µ=95.3% (σ=2.5%) with µ=99.6% (σ=1.2%) correctly decoded trials in the *attention* class and µ=73.8% (σ=16.7%) correctly decoded trials in the *passive viewing* class. In experiment 2 the CV revealed a decoding accuracy of µ=93.5% (σ=2.5%) with µ=99.8% (σ=0.7%) correctly decoded trials in the *attention* class and µ=62.2% (σ=13.4%) correctly decoded trials in the *passive viewing* class. The high sensitivity and comparably low specificity indicate that the decoder is biased towards the *attention* class with more training samples available. Increasing the number of training samples in the *passive viewing* class will most likely increase the specificity.

### 3.4. Effects of visual spatial attention

The effects of visual spatial attention, essentially driving the decoder we analysed with cluster-based permutation tests as well. We compared attention to targets presented in the right visual field (RVF) with attention to targets presented in the left visual field (LVF). In experiment 1, an early parieto-occipital cluster of 10 left electrode sites (z_clust_=-303, p_clust_=0.0619) showed a stronger deflection to right vs left targets between 164ms to 230ms following stimulus onset (peak difference at 197ms at electrode PO3; p_sr_ =.0003; see Figure 6A). A similar cluster was found in experiment 2, where a cluster of 11 electrode sites (z_clust_=-304, p_clust_=0.0639) showed a significant difference between LVF and RVF, ranging from 156ms to 230ms following stimulus onset (peak difference at 197ms at electrode PO3; p_sr_ =.0002; see Figure 6B). These clusters did not meet the significance criterion of p_clust_<0.05 but the p-value at the difference peak clearly indicated a significant difference. Their temporal and spatial characteristics indicate that the observed effects correspond to the N2pc component (Luck and Hillyard, 1994). A second occipital-temporal cluster of 10 left electrodes (z_clust_=-430, p_clust_=0.0220) showed a stronger negative deflection to contralateral targets between 271ms to 477ms (peak difference at 386ms at P7, p_sr_ =.0005) in experiment 1. Similarly, a cluster of 9 electrode sites (z_clust_=-734, p_clust_=0.0100) showed a stronger deflection to LVF vs RVF between 288ms and 600ms ms following stimulus onset (peak difference at 403ms at electrode P7; p_sr_ =.0004) in experiment 2. This late extended component is described in the literature as sustained posterior contralateral negativity (SPCN) (Jolicœur et al., 2008), typically following the N2pc.

**Figure 6:**
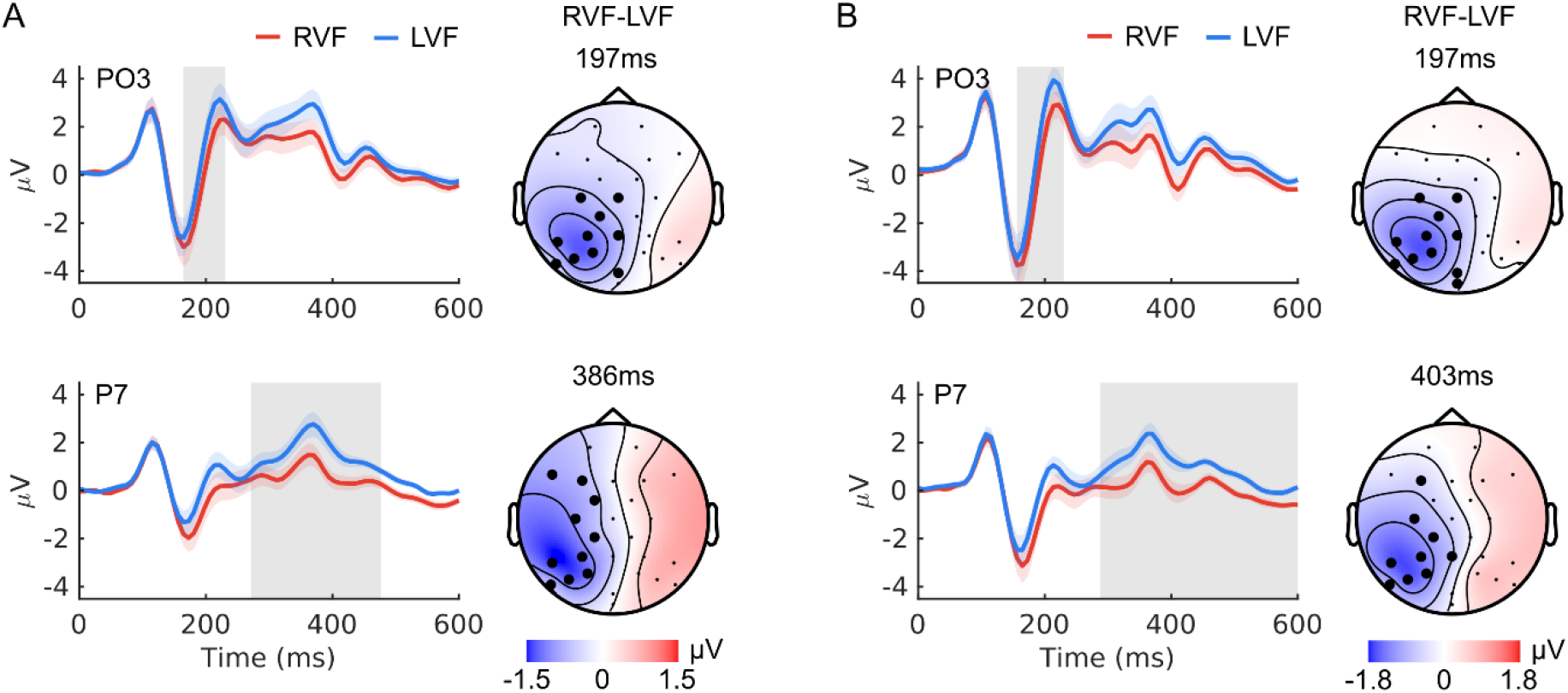
Visual spatial attention effects. Diagrams show the average signals for attention to targets in the right visual field (RVF) and left visual field (LVF) at the electrode with greatest difference in a cluster. Grey areas indicate the range of significant cluster time points at that electrode. Enlarged electrode markers indicate cluster channels at the time point of greatest difference in **(A)** experiment 1 and **(B)** experiment 2.

### 3.5. Effects of eye movements

Although participants were instructed to keep their gaze fixed while paying attention to peripheral stimuli, unintended saccades might have been performed. We trained a decoder model similarly as with the EEG but using the recorded EOG. The average decoding accuracy using all attention trials was µ=66.1% (σ=7.7%) for experiment 1 and µ=64.8 (σ=8.7%) for experiment 2. EOG decoding of 8 participants did not exceed the guessing level of 60.1% as obtained by permutation testing. The EOG-based decoding accuracies of all participants were numerically smaller than the EEG-based decoding accuracies (p<0.001). The correlation of EOG and EEG-based decoding accuracies across participants was not significant (r=0.29, p=0.24 for experiment 1 and r=0.12, p=0.65, for experiment 2).

We also determined the deflection of the horizontal EOG following each visual stimulus. On average, the deflection of the horizontal EOG was µ=8.69 µV (σ=4.86 µV) during attention trials and µ=6.74 µV (σ=1.35 µV) during ignored trials in experiment 1. This difference was statistically significant (p=0.008). In experiment 2, the average deflection of the horizontal EOG was also higher during attention trials (µ=7.4 µV, σ=1.79 µV) compared to ignored trials (µ=6.88 µV, σ=1.37 µV), but the difference was not statistically significant. In both experiments, we found no significant difference between the deflection values, neither between binocular, right-eye and left-eye vision nor between normal, moderately degraded and highly degraded vision.

## 4. Discussion

In this study we systematically investigated the impact of different viewing conditions on event-related potentials and on decoding of covert attention to control a BCI. While we found spatio-temporal clusters with differences in the event-related potentials between viewing conditions indicating impaired visual processing, the decoding accuracy, which is an indicator for BCI performance, was not altered.

BCIs that rely on visual stimulation have been shown to provide reliable signals to decode a user’s intention from event-related potentials related to attention. However, vision-based BCIs have the disadvantage that visual acuity declines with increasing distance from the fovea (Anstis, 1974). This makes discrimination of peripheral items without gaze shifts a challenge and considerably reduces BCI performance (Brunner et al., 2010). Importantly, BCIs developed and tested with healthy participants usually do not take vision impairments into account. Here, healthy participants performed a VSA task while different visual impairments were simulated. In the first experiment, we covered one eye of participants to compare monocular vison and binocular vision. In the second experiment, visual acuity was reduced by using Bangerter occlusion foils, which reduces visual acuity to a specific value (e.g. 0.1 visual acuity equals Snellen index 6/60). The gaze-independent BCI protocol we used evokes reliable attention-based EEG components at the chosen eccentricity and size of stimuli (Reichert et al., 2020b). Furthermore, color alone as the target feature allows high decoding accuracies larger than 90% correct (Reichert et al., 2022), making the paradigm well suited for users with impaired vision. Comparable with these studies we achieved decoding results > 90% independently of impaired vision. This contradicts the assumption that vision impairment, which has been reported to modulate EEG responses (Collins et al., 1979), affects VSA-based BCI performance. Because correlation is a scale-invariant measure, amplitude differences between canonical ERPs do not influence classification in CCA-based decoding. Instead, the decoder relies on the temporal sequence of attentional allocation reflected in the evolution of the ERP over time. Furthermore, canonical components were robust when test and training data were separated according to the conditions. Only monocular right-eye vision components predicted the monocular left-eye vision with reduced decoding accuracy due to opposing lateralization of left and right monocular vision compared to binocular vision. Similarly, moderately degraded vision was predicted with reduced accuracy by canonical components obtained from the other viewing conditions, which is in line with different spatial patterns in the significant clusters we found. However, the reduction in decoding accuracy was purely numerical and does not undermine the overall robustness of the decoder.

Furthermore, in both experiments we found overlapping spatial clusters showing the difference between left and right presented targets. The clusters showed modulation differences in the N1 and P2, which is in line with previous studies (Hopf et al., 2000; Jolicœur et al., 2008; Luck and Hillyard, 1994; Woodman and Luck, 1999). The N2pc component has been shown to be independent of probability as well as of the feature dimensions orientation, color, and size (Luck and Hillyard, 1994). Commonly, the N2pc and SPCN are calculated from collapsed signals of both hemispheres to increase the signal-to-noise ratio and focus on contralateral effects. Although this is a standard approach in vision research literature, it neglects hemispheric differences. However, since these differences are advantageous for a decoder relying on spatial filtering, we did not collapse the signals of both hemispheres. Thus, we compared left and right target presentation directly in the statistical analysis rather than comparing contralateral and ipsilateral presentation. This resulted in clusters in the left but not in the right hemisphere indexing both N2pc and SPCN activity. Such hemispheric dominance has also been observed in related visual search studies (Eimer, 1996; Reichert et al., 2020a). Comparing monocular and binocular vision independent of target location, we found three clusters. The steeper P1 and N1 components during binocular vision compared to monocular vision indicates that monocular vision results in delayed sensory processing compared to binocular vision, which is in line with results obtained from checkerboard reversal stimuli (Adachi-Usami and Lehmann, 1983; di Summa et al., 1997). The spatial distribution of N1 differences contralateral to the covered eye can be explained by the binocular advantage, which results from binocular summation (Baker et al., 2018) and implies faster reaction times (Wakayama et al., 2011). Paramacular N1 generators are especially sensitive to interocular asymmetries (Jeffreys and Axford, 1972), such that removing input from one eye slows the N1 response in the contralateral hemisphere. In contrast, the P2 component, assumed to be involved in attention processes (Di Russo et al., 2012; Treder and Blankertz, 2010), showed maximal differences at the parietal midline because its extrastriate generators integrate signals bilaterally and are less sensitive to differences in monocular contribution (Shawkat and Kriss, 1997).

The P3 component is considered an electrophysiological marker of the detection and evaluation of target stimuli (Polich, 2007) and is elicited when a stimulus is categorized as task-relevant (Luck, 2014). In the current study, clusters indexing P3 modulation have been found in several statistical tests. Comparison of normal and degraded vision resulted in a significant cluster indicating that the P3 amplitude was increased with normal vision compared to highly degraded vision. MDV showed increased P3 compared to HDV as well, where the localization of the maximal difference was at Cz. In contrast, Nv and HDV most deviated at bilateral parieto-occipital electrodes. The reduced P3 amplitude under HDV may reflect participantś inability to recognize the letter presented in the target color. Finally, our investigation on attention epochs in comparison to passive viewing epochs revealed a clear P3 pattern typically found in attention experiments and often used for BCI control in oddball paradigms. The fact that attention can be reliably decoded from this pattern, which was demonstrated by our decoding approach achieving high sensitivity rates, opens the possibility to use the attention/passive-viewing decoder as a switch to determine communication intention at all in a BCI setting that otherwise would expect a response on each trial.

Our decoding results and the results from the statistical analysis demonstrate that impairment of vision can be compensated for by attention processes as long as the target features can be distinguished. Since the target feature color can be distinguished by persons with impaired vision, our BCI protocol produced equally reliable results in all stages of visual impairment we tested, making it suitable for people suffering from visual blur, diplopia and ophthalmoplegia. It therefore represents an alternative approach to auditory and motor-imagery protocols for gaze-independent BCI control in the visual domain.

## Ethics approval statement

This study was conducted in accordance with the Declaration of Helsinki and was approved by the Ethics Committee of the Medical Faculty and the University Hospital of the Otto von Guericke University, Magdeburg (No. 158/22).

## CRediT authorship contribution statement

**Christoph Reichert:** Conceptualization, Investigation, Funding acquisition, Software, Visualization, Writing – Original Draft. **Mircea Ariel Schoenfeld:** Formal Analysis, Resources, Validation, Writing – review and editing. **Stefan Dürschmid:** Conceptualization, Project administration, Supervision, Validation, Writing – Review & Editing.

## Acknowledgments

The authors thank Julie Morgan for assisting in the data collection.

